# Multimodal Hox5 activity generates motor neuron diversity

**DOI:** 10.1101/2024.02.08.579338

**Authors:** Ritesh KC, Raquel López de Boer, Minshan Lin, Lucie Jeannotte, Polyxeni Philippidou

## Abstract

Motor neurons (MNs) are the final output of circuits driving fundamental behaviors, such as respiration and locomotion. Hox proteins are essential in generating the MN diversity required for accomplishing these functions, but the transcriptional mechanisms that enable Hox paralogs to assign distinct MN subtype identities despite their promiscuous DNA binding motif are not well understood. Here we show that Hoxa5 controls chromatin accessibility in all mouse spinal cervical MN subtypes and engages TALE co-factors to directly bind and regulate subtype-specific genes. We identify a paralog-specific interaction of Hoxa5 with the phrenic MN-specific transcription factor Scip and show that heterologous expression of Hoxa5 and Scip is sufficient to suppress limb-innervating MN identity. We also demonstrate that phrenic MN identity is stable after Hoxa5 downregulation and identify Klf proteins as potential regulators of phrenic MN maintenance. Our data identify multiple modes of Hoxa5 action that converge to induce and maintain MN identity.

## Introduction

The motor programs that mediate essential behaviors such as respiration and locomotion rely on the establishment of distinct subtypes of motor neurons (MNs) during development. MN diversity arises from the intersection of dorsoventral and rostrocaudal signaling pathways that drive the combinatorial expression of unique sets of transcription factors (TFs) that specify MN subtype identities along the spinal cord^1,2^. Along the rostrocaudal axis, members of the chromosomally-clustered *Hox* gene family are critical in specifying the identity of segmentally-restricted MN subtypes^3^. Despite the well-described functions of Hox proteins in MN specification, several questions remain regarding the mechanisms that different Hox paralogs employ to induce distinct subtype identities at the transcriptional level and how Hox protein divergent and convergent functions are mediated^4,5^. For example, while several Hox proteins have been shown to converge on common transcriptional targets to redundantly promote limb-innervating Lateral Motor Column (LMC) identity^6^, it is less clear how a single Hox paralog may promote multiple MN subtype identities.

Hox proteins bind DNA through their homeodomain, a 60 amino acid domain that recognizes a short TA-rich DNA motif. Homeobox domains are highly similar amongst different Hox proteins and do not appear to confer DNA-binding selectivity to individual paralogs^7–10^. This contrasts with the unique functions of Hox proteins in vivo, which implies a stringent selectivity of gene targets, giving rise to the Hox-specificity paradox^11,12^. How do Hox proteins achieve their unique functions given the apparent overlap in their DNA-binding motifs? One partial solution to this paradox arises from the cooperative binding of Hox proteins to DNA with a family of cofactors, known as the three aminoacid loop extension (TALE) homeodomain proteins^13^. Pbx proteins, members of the TALE family of TFs, are essential mediators of Hox function in MNs and mutations in Pbx genes recapitulate Hox mutant phenotypes^14^. While Hox/Pbx interactions increase the specificity of the DNA-binding site, it is unlikely that this interaction alone accounts for all the unique functions of individual Hox paralogs, as multiple Hox proteins are able to interact with Pbx proteins, pointing to the existence of additional mechanisms that further contribute to Hox specificity^15^.

At cervical levels of the spinal cord, Hox5 paralogs have the ability to promote both Phrenic Motor Column (PMC) and LMC identity^6,16^. Mice lacking *Hox5* genes in MNs die at birth from respiratory failure, largely due to progressive loss and disorganization of phrenic MNs, and a dramatic loss in axon branching and synaptic contacts at the diaphragm^16^. Effects on limb-innervating MNs are subtler, as *Hox5* mutant mice show grossly normal patterns of limb innervation, with only a subset of motor pools adopting abnormal trajectories and targeting inappropriate muscles^17^. The transcriptional mechanisms that underlie the ability of a single Hox TF to induce two opposing MN identities are not well understood. Hox5 proteins are the only Hox paralogs that induce PMC-specific genes in vivo, while the ability to induce genes expressed in LMC neurons is common with other Hox family members (Hox4-8). How do Hox5 proteins accomplish both unique and shared functions in MNs? One possibility is that this distinction arises through different DNA-binding motifs which are highly Hox5-specific in PMC genes but common for multiple Hox proteins in LMC genes. An example of this can be seen in *Drosophila*, where the Hox5 homolog Sex combs reduced (Scr), the only Hox protein that can initiate salivary gland development, can bind cooperatively with the Pbx homolog Extradenticle (Exd) to a unique sequence that other Hox/Pbx complexes are unable to bind^18^. Do Hox5 proteins act in a similar manner in phrenic MNs to bind Hox5/Pbx specific sites? While this mechanism of action might account for the unique ability of Hox5 proteins to induce PMC-specific genes, it would fail to explain how these genes are restricted specifically to the PMC given the co-expression of Hox5 and Pbx proteins in other MN populations in the cervical spinal cord. An alternative hypothesis is that additional DNA-binding proteins contribute to the selection of specific targets, either by forming a complex with Hox5/Pbx and altering the preference for a binding site, or by differentially recruiting activators or repressors to the transcriptional complex. In addition to Hox5 and Pbx proteins, PMC neurons also express the POU-domain transcription factor (TF) Scip (Pou3f1, Oct6)^16,19,20^ while LMC neurons express the TF FoxP1, which is required for the induction of Hox-dependent LMC-specific genes^21,22^. Therefore, one possibility is that, depending on the presence of either Scip or FoxP1, Hox5/Pbx/Scip and Hox5/Pbx/FoxP1 complexes activate two non-overlapping sets of targets, required for PMC and LMC specification respectively.

In addition to their canonical functions as TFs, Hox paralog activities can also diverge based on their differential ability to open chromatin, a characteristic property of pioneer factors^23–25^. For example, Hox13 pioneer activity is essential for initiating developmental programs required for the generation of limb digits and external genitalia in mammals^26,27^. During in vitro MN specification, Hox TFs exhibit differential abilities to bind and open inaccessible chromatin^28^. Hox5 proteins may partly act by promoting the opening of chromatin that is actively-transcribed in specific MN columns. The ability of Hox proteins to alter chromatin state might also contribute to the stable maintenance of subtype-specific MN identity after the downregulation of Hox proteins at postnatal stages.

Here, we utilize Assay for Transposase-Accessible Chromatin using sequencing (ATAC-seq) from isolated mouse embryonic MNs to show that Hox5 TFs possess the ability to open chromatin associated with all three major columns of MNs in the cervical spinal cord and engage TALE co-factors to directly bind and regulate subtype-specific genes. We identify a paralog-specific interaction of Hoxa5 with Scip and show that heterologous expression of Hoxa5 and Scip is sufficient to suppress alternative MN identities. We also demonstrate that phrenic MN identity is stable after Hox5 downregulation and identify Klf TFs as potential downstream regulators of phrenic MN maintenance. Our data identify multiple modes of Hox5 action that converge to induce and maintain MN identity.

## Results

### Hoxa5 regulates cervical MN chromatin accessibility

Spinal MNs are generated from a highly restricted common progenitor domain in the ventral neural tube. As MNs begin to differentiate and exit the cell cycle, they are topographically organized in a stereotypical fashion as discrete motor columns which exhibit distinct transcriptional profiles and subtype-specific molecular markers by embryonic day (e)12.5. Cervical levels of the spinal cord contain MNs that can be divided into three major subtypes: Phrenic Motor Column (PMC) neurons which innervate the diaphragm to regulate breathing, Lateral Motor Column (LMC) neurons that project to the upper limbs, and Medial Motor Column (MMC) neurons that project to dorsal axial muscles to control posture (Fig. 1a).

**Figure 1.**
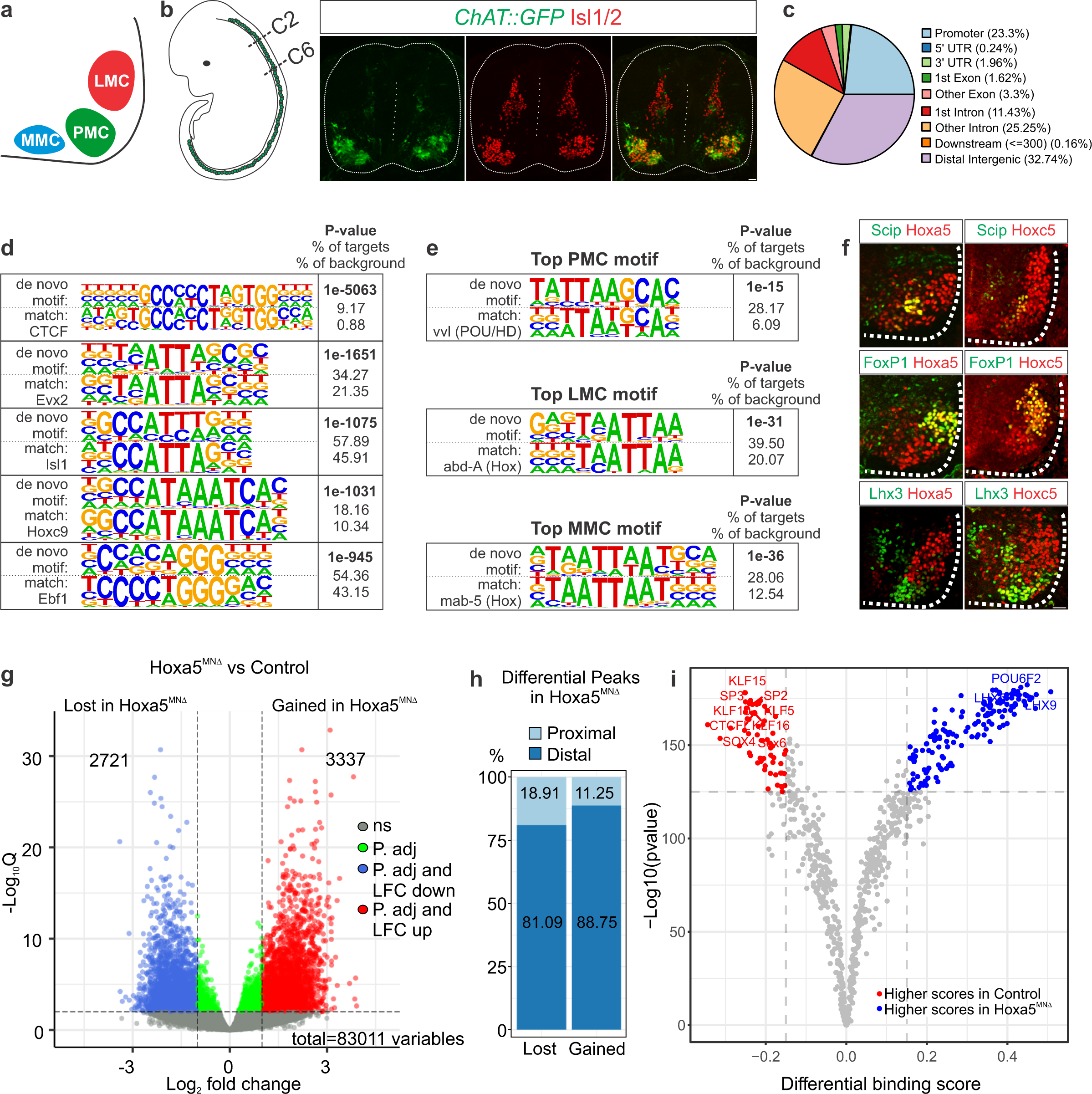
Hoxa5 contributes to chromatin accessibility in cervical motor neuron (MN) subtypes. **a)** MN subtypes at cervical levels of the spinal cord. Phrenic Motor Column (PMC) neurons innervate the diaphragm, Lateral Motor Column (LMC) neurons project to limb muscles, and Medial Motor Column (MMC) neurons innervate axial muscles. **b)** *ChAT:: GFP* reporter mice label MNs in green, as seen by co-localization with the MN-specific transcription factor (TF) Isl1/2 (red). We used fluorescence activated cell sorting (FACS) to isolate GFP+ MNs from spinal cervical levels C2-C6 for Assay for Transposase-Accessible Chromatin using sequencing (ATAC-seq) analyses. Scale bar= 50mm. **c)** Distribution of ATAC-seq peak location relative to the nearest transcription start site (TSS). **d)** HOMER output of top motifs enriched in ATAC-seq peaks. Both the de novo motif (top) and the best matched known TF motif (bottom) are shown, along with p-value and prevalence. **e)** Top HOMER motif identified for motor column-specific genes after either intersection with scRNA-seq data (LMC and MMC) or examination of known column-specific genes (PMC). De novo motifs match known Hox motifs for all columns. **f)** Hox5 paralog expression in cervical motor columns. Both PMC (Scip+) and LMC (FoxP1+) neurons show high expression levels of Hoxa5 and Hoxc5. MMC (Lhx3+) neurons express low levels of both Hoxa5 and Hoxc5, while Hoxb5 is not expressed in MNs (Fig. S1c). Scale bar= 100mm. **g)** Volcano plot showing differential chromatin accessibility between control and *Hoxa5^MNΔ^* MNs, determined by DESeq2, with fold change cutoff of 2-fold and significance cutoff of FDR < 0.01. 3337 peaks were significantly gained, while 2721 peaks were significantly lost in *Hoxa5^MNΔ^* MNs. **h)** Distribution of differential ATAC-seq peaks in *Hoxa5^MNΔ^* MNs. **i)** Comparison of TF activities between control and *Hoxa5^MNΔ^* MNs. Volcano plot showing the TOBIAS differential binding score on the x-axis and -log10(p value) on the y-axis; each dot represents one TF.

To gain insights into the transcriptional programs that regulate MN specification and diversity, we performed ATAC-seq to identify regions of actively-transcribed open chromatin in cervical MNs. We used *Choline Acetyltransferase (ChAT)::GFP* transgenic reporter mice, which express GFP in ventrally located Isl1/2+ MNs, to sort MNs from the cervical spinal cord at e12.5, when motor columns have acquired their distinct identities (Fig. 1b, S1a). We generated ATAC-seq biological replicates with a mean of 107M unique paired-end mapped reads per sample and identified 85,866 peaks of transposase accessible chromatin that were distributed across both intronic and exonic regions, with about 23% being located in promoters (1kb upstream or downstream of the transcriptional start site (TSS), including peaks at the *ChAT* promoter and the pan-MN TFs, *Isl1* and *Mnx1 (Hb9)* (Fig. 1c, S1b). Next, we used HOMER^29^ to perform de novo motif search using ATAC-seq peaks to find the relative abundance of sequence-specific TF consensus motifs. We identified the enrichment of CTCF motifs along with known MN markers such as Isl1 and Ebf1, as well as prominent homeobox recognition motifs such as Evx2 and Hoxc9 (Fig. 1d).

To identify chromatin accessibility regions that correspond to distinct MN columnar subtypes (PMC, LMC and MMC), we compared our ATAC-seq generated peaks to column-enriched genes identified by scRNA-seq^30^. We employed a graph-based clustering approach, Seurat^31^, to identify the expression of differential genes. We assigned columnar identities based on the average expression of key MN marker genes in distinct clusters. For example, MN clusters exhibiting high expression of Foxp1 and Aldh1a2 were combined and assigned as LMC. Similarly, MNs exhibiting high expression of Mecom and Lhx3 were combined and assigned as MMC. With this approach, however, we were unable to confidently identify a phrenic MN cluster, likely due to the fact that PMC neurons are a rare population that may not form a distinct cluster in embryonic scRNA-seq data. Therefore, we instead utilized a list of genes known to be selectively enriched in phrenic MNs by in situ hybridization^19,32^. We then assigned ATAC-seq peaks to the gene of their nearest TSS and intersected genes associated with ATAC-seq peaks with column-enriched genes (Table S1).

To identify unique regulators of each MN subtype we performed motif analysis restricted to column-enriched genes. Our analysis identified prominent Hox motifs in the regulatory regions of MNs belonging to all three columns, indicating the predominant role of Hox TFs in MN specification (Fig. 1e). At cervical levels of the spinal cord, Hox5 paralogs (Hoxa5, Hoxb5 and Hoxc5) are the major Hox proteins expressed^33^. While Hox5 proteins have been implicated in both PMC and LMC development^6,16,17,33^, we were surprised to identify Hox motifs in MMC neuron-enriched genes, as MMC development is thought to be Hox-independent^21,22^. We confirmed that both PMC and LMC neurons express high levels of Hoxa5 and Hoxc5, while also detecting Hoxa5 and Hoxc5 expression in MMC neurons at lower levels (Fig. 1f)^16^. Hoxb5 is not expressed in MN populations at e12.5 (Fig. S1c). To test whether Hoxa5 regulates chromatin accessibility, we performed ATAC-seq on sorted *Hoxa5*-deleted (*Hoxa5flox/flox; Olig2::Cre*, referred to as *Hoxa5^MNΔ^*) cervical MNs. Principal component analysis (PCA) showed a high degree of concordance between replicates, with *Hoxa5* deletion accounting for the majority of variance (Fig. S1d).

To define chromatin accessibility changes induced by the loss of *Hoxa5*, we performed differential analysis using DESeq2. We identified a total of 3337 and 2721 peaks that were either gained or lost in Hoxa5^MNΔ^ MNs, respectively (q-value <0.01, ±2x) (Fig.1g). To test whether Hoxa5 differentially alters chromatin accessibility at promoter or enhancer regions, we analyzed the distribution of differential ATAC-seq peaks at proximal (≤2000kb) and distal (>2000kb) regions from an annotated TSS. While the majority of peaks that are gained or lost in *Hoxa5^MNΔ^* MNs are distributed at distal enhancers, there is a higher percentage of peaks with decreased accessibility in *Hoxa5^MNΔ^* MNs located at promoter regions, suggesting that Hoxa5 may have a different impact on chromatin accessibility at proximal and distal regulatory elements (Fig. 1h).

To identify molecular pathways impacted by chromatin changes after Hoxa5 loss, we performed GO term enrichment analysis using the nearest annotated neighboring genes for individual chromatin accessibility peaks. We found that genes with altered accessibility are associated with developmental processes such as axonogenesis, neurogenesis, regionalization, axon guidance, dendrite development, synapse organization and cell adhesion, consistent with the phenotypes observed in Hox5^MNΔ^ mice (Fig. S1e)^16,32^. Altogether, these results suggest that Hox5 TFs regulate MN-specific gene expression programs partly by altering the MN chromatin landscape.

To define the TFs that are enriched in differentially accessible regions and thus may, in addition to Hoxa5, control MN gene regulatory programs, we performed footprinting analysis in control and Hoxa5^MNΔ^ ATAC-seq peaks, using TOBIAS^34^ with motifs from the Jasper databases^35^. This computational approach uses transposase insertion sites to identify motifs that are protected from transposition, hence likely bound by a TF. Differential footprinting analysis showed that motifs for Klf TFs (Klf5, Klf15, Klf10) showed a higher footprinting score in control peaks, whereas motifs for homeobox TFs such as Lhx showed a higher footprinting score in Hoxa5^MNΔ^ peaks. The high occurrence of Klf motifs in control peaks suggests that Hoxa5-mediated chromatin reorganization may expose previously-inaccessible Klf binding sites. Overall, the differential footprints found between control and Hoxa5^MNΔ^ MNs support the idea that Hoxa5 may regulate the binding ability of downstream TFs.

### Hoxa5 and Pbx1 modules directly control MN genes

To understand how Hoxa5 induces distinct MN subtype identities, we wanted to identify direct Hoxa5 transcriptional targets in the spinal cord. Since TALE cofactors cooperatively bind chromatin with Hox proteins and are essential for many Hox actions, we also investigated targets of Pbx1, which is strongly expressed in all cervical MN columns^14^. To identify both unique and shared transcriptional targets of Hoxa5 and Pbx1, we performed chromatin immunoprecipitation followed by sequencing (ChIP-seq) from e12.5 mouse cervical spinal cord chromatin, and identified a total of 3494 Hoxa5 peaks and 15764 Pbx1 peaks. To understand if Hoxa5 and Pbx1 co-regulate a subset of cis regulatory elements, we intersected Hoxa5 with Pbx1 peaks and found that 34% of Hoxa5 (1186) peaks co-occur with Pbx1 peaks (Fig. 2g). The majority of Pbx1 peaks were not bound by Hoxa5, indicating that many Pbx functions are likely Hoxa5-independent^36^. Notably, we also identified a significant portion of Hoxa5 peaks not bound by Pbx1 indicating either that other Pbx proteins, such as Pbx3, may form distinct complexes with Hoxa5, or suggesting Pbx-independent Hoxa5 DNA binding. Further analysis of the genomic distribution of Hoxa5 and Pbx1 bound regions showed that the majority of Hoxa5 peaks were located within the promoter region of annotated genes, while Pbx1 peaks were distributed between promoters, intronic and intergenic regions (Fig. 2a-b, 2d-e). Regions co-occupied by both Hoxa5 and Pbx1 are predominantly associated with promoter regions, mostly mirroring Hoxa5 peak distribution (Fig. 2h). GO term enrichment analysis revealed that the peaks bound by either Hoxa5 or Pbx1 or both are associated with genes that regulate axonogenesis, pattern specification process, regionalization, and migration of neurons (Fig. 2k, S2a-b), consistent with known Hoxa5/Pbx1 functions in MNs. To investigate whether certain DNA motifs were enriched in the Hoxa5-bound, Pbx1-bound, and combined Hoxa5-Pbx1 bound sites, we applied HOMER de novo motif search. Surprisingly, the top motifs identified in Hoxa5-bound sites were not canonical Hox motifs, but were instead enriched for Neurod1 and E2F2 consensus binding sequences, while Pbx1-bound sites were enriched for Hox, Pbx and Meis motifs. A previously established Hox-Pbx composite motif (Fig. 2j) was identified in the top enriched motifs in all Hoxa5-bound, Pbx1-bound, and Hoxa5-Pbx1 shared peaks (Fig 2c, 2f, 2i).

**Figure 2.**
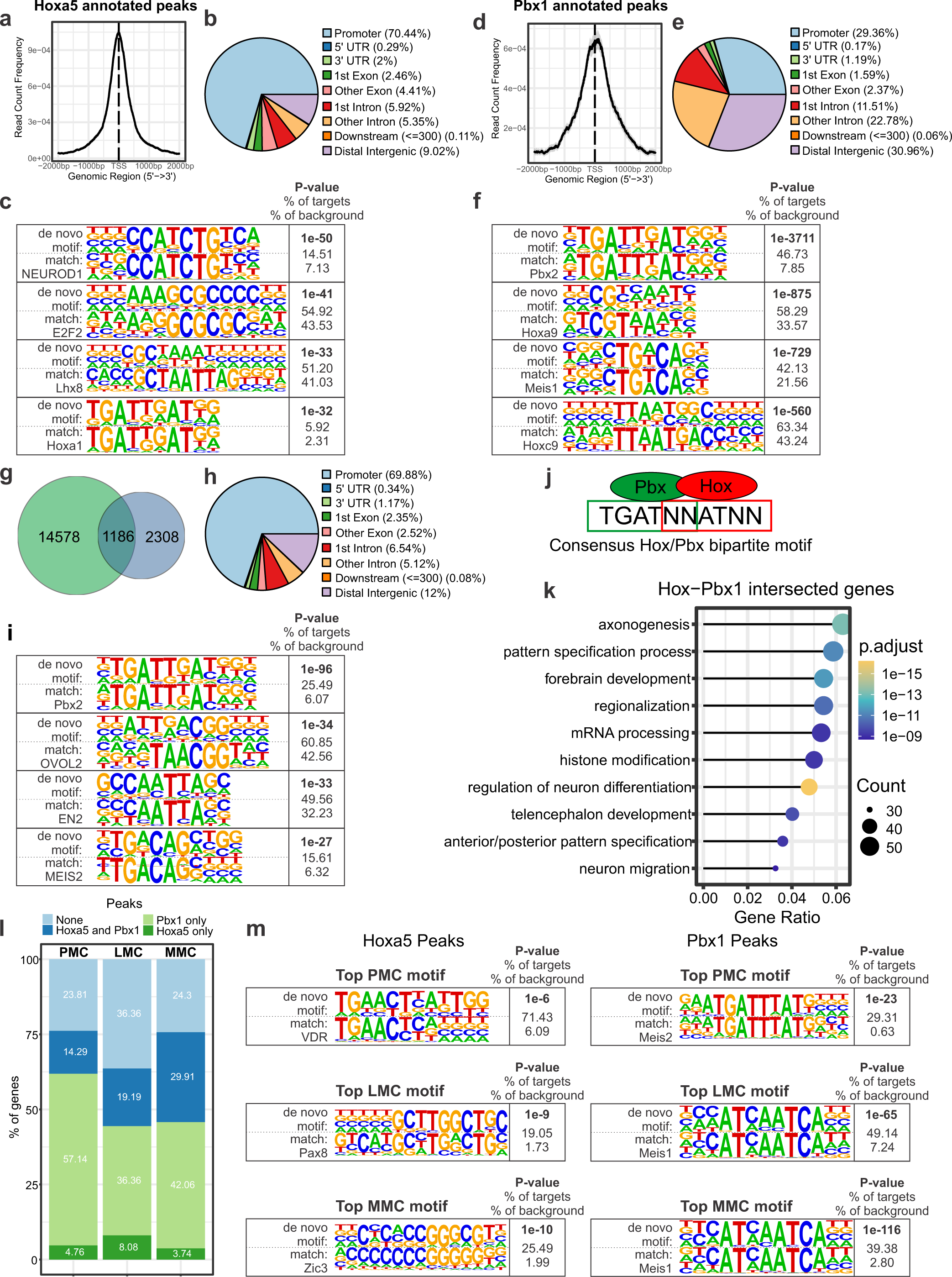
Direct regulation of MN-specific genes by Hoxa5 and Pbx1. **a, d)** Average distribution around the TSS of Hoxa5 **(a)** and Pbx1 **(d)** target genes. **b, e)** Pie chart illustrating peak location relative to the nearest TSS for Hoxa5 **(b)** and Pbx1 **(e)** enriched peaks. The distribution of Hoxa5-bound peaks is enriched in promoters compared to Pbx1. **c, f)** HOMER output of top motifs enriched in Hoxa5 **(c)** and Pbx1 **(f)**-bound peaks. **g)** Overlap of Hoxa5 and Pbx1 enriched peaks. **h)** Pie chart illustrating peak location relative to the nearest TSS for Hoxa5 and Pbx1 enriched peaks. **i)** HOMER output of top motifs enriched in Hoxa5- and Pbx1-bound peaks. **j)** Consensus Hox/Pbx bipartite motif. **k)** Gene ontology (GO) enrichment analysis of biological pathways of genes with Hoxa5-Pbx1 intersected ChIP-seq peaks. Top 10 significant GO enrichment pathways shown based on the gene counts in each category. **l)** Analysis of column-specific genes for Hoxa5 and Pbx1 binding. **m)** Top HOMER motif identified for

Next, we associated Hoxa5, Pbx1 and Hoxa5-Pbx1 intersected peaks to the major MN column genes (Fig. 2l). Our analysis identified that more than 75% of PMC and MMC genes show enrichment of either Hoxa5 or Pbx1 or both, underscoring the overarching function of Hox and Pbx-mediated transcriptional programs in these MN populations. In contrast, we found that a significant portion of LMC genes (36%) do not show enrichment of either Hoxa5 or Pbx1, likely reflecting Hox-downstream programs that regulate MN pool identity. Assessment of TF motifs present in Hoxa5-bound PMC, LMC, and MMC genes using HOMER revealed enrichment of distinct motifs for each column, indicating that the specific cellular context in each MN subtype might alter Hoxa5 binding specificity (Fig. 2m). De novo motif analysis of Pbx1-bound peaks in all motor columns revealed enrichment of motifs for the TALE cofactors Meis1 and Meis2. This suggests that Pbx1 may bind specific motor column loci in a Hoxa5-independent manner, in a complex with other TALE factors such as Meis1 and Meis2. Together, these results suggest that Hoxa5 and Pbx1 either individually or collaboratively target cis-regulatory modules that orchestrate different aspects of MN development.

### Scip cooperates with Hox/Pbx programs to induce PMC identity

We found that Hoxa5 and Pbx1 directly bind and regulate genes that are essential for MN specification and development in multiple motor columns in the cervical spinal cord. However, it is unclear how these TFs can induce specific MN identities given their broad expression pattern. We previously showed by retrograde labeling that the expression of Scip, a POU domain TF, is restricted to PMC neurons and that overexpression of FoxP1, which is required for the establishment of LMC identity^21,22^, suppresses Scip expression^16^. We next asked whether context-specific functions of Hoxa5 are achieved via interactions or cooperativity with other MN-specific TFs, such as Scip.

To test whether Hoxa5 and Scip associate with each other during PMC specification, we created tagged constructs (Fig. 3a) and performed protein co-immunoprecipitation (co-IP) assays using transiently transfected 293T cells. As a control experiment, we also looked at the interaction between Hoxa5 and Pbx1, which has been previously established^37,38^. The co-elution of Hoxa5 and Pbx1 and Hoxa5 and Scip in the same IP fraction, suggests that these proteins can form a complex (Fig. 3b, c). The hexapeptide (YPWM) domain of Hox proteins is critical for their interaction with Pbx cofactors^37,39,40^. To test whether the same domain is required for the Hoxa5 interaction with Scip, we mutated the YPWM domain of Hoxa5 to AAAA (Hoxa5^YPWM>AAAA^, fig.3a) and performed co-IP assays. While we found a decreased association of Hoxa5^YPWM>AAAA^ with Pbx1 as expected, we did not find any changes in its interaction with Scip, suggesting that the Hoxa5-Scip interaction is independent of the hexapeptide motif. These data support a model where Hoxa5, Pbx1, and Scip form a complex to induce phrenic-specific programs and both Pbx1 and Scip bind to Hoxa5 through non-competitive interactions.

**Figure 3.**
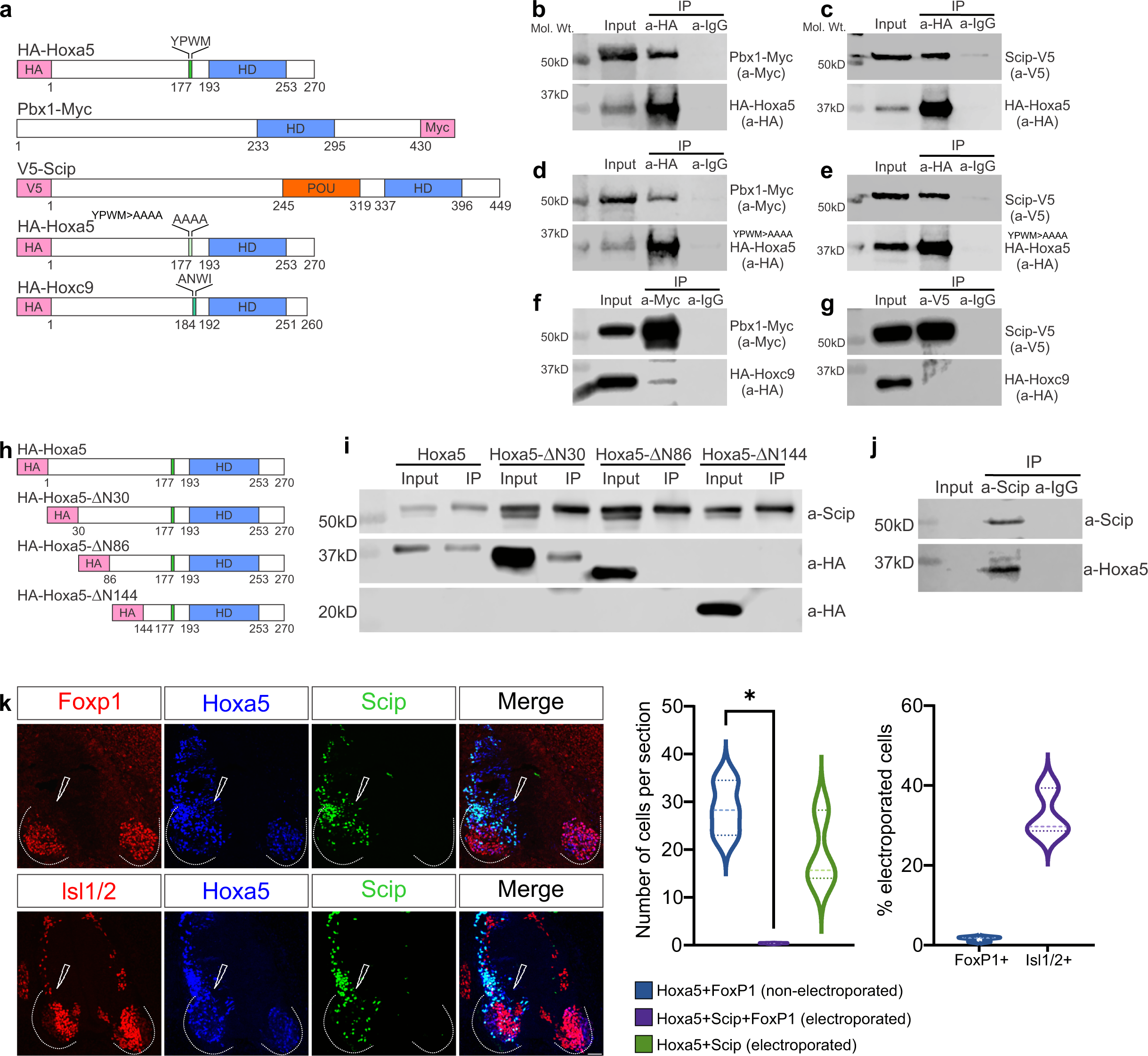
Paralog-specific Hoxa5/Scip interaction promotes PMC identity. **a)** Overview of tagged constructs used for co-immunoprecipitation (co-IP) in transiently transfected 293T cells. **b-c)** HA-Hoxa5 co-immunoprecipitates with Pbx1-Myc and V5-Scip. **d-e)** HA-Hoxa5^YPWM>AAAA^ co-immunoprecipitates with V5-Scip **(e)** while co-immunoprecipitation with Pbx1-Myc **(d)** is reduced. **f-g)** HA-Hoxc9 does not interact with V5-Scip **(g)** and weakly interacts with Pbx1-Myc **(f)**. **h)** Overview of Hoxa5 N-terminal serial truncation constructs. **i)** Transiently transfected 293T cells with HA-Hoxa5 N-terminal serial deletion constructs and Scip-V5 were subjected to co-immunoprecipitation assay using antibodies against V5. Hoxa5 and ΔN30 Hoxa5 co-immunoprecipitate with V5-Scip, while Hoxa5-NΔ86 and Hoxa5-NΔ144 do not. **j)** Scip and Hoxa5 co-immunoprecipitation from whole cell lysate of e12.5 embryonic spinal cord tissue. **k)** Co-electroporation of Hoxa5 and Scip in chick embryos leads to a reduction in the number of FoxP1 positive cells in the cervical spinal cord, but does not affect overall MN identity, as seen by Isl1/2 expression (n=3), p= 0.0256. Scale bar= 50μm.

To understand whether the Hoxa5 interaction with Scip is paralog-specific, we tested the ability of Scip to interact with Hoxc9, a Hox paralog required for the generation of thoracic respiratory MN subtypes that is ∼36% identical to Hoxa5^41^. We found that Hoxc9 does not form a complex with Scip (Fig. 3g), suggesting that Scip does not broadly associate with Hox proteins, but rather exhibits paralog-dependent specificity. Due to the absence of a canonical hexapeptide motif, Hoxc9 also shows decreased interaction with Pbx1 (Fig. 3f)^40^.

Outside of the homeodomain and the YPWM motif, N-terminal domains of Hox protein sequences diverge substantially. To identify the region of Hoxa5 necessary for complex formation with Scip, we serially deleted the N-terminal end of Hoxa5 and created three HA-tagged N-terminal deletion constructs: HA-Hoxa5-ΔN30, HA-Hoxa5-ΔN86 and HA-Hoxa5-ΔN144 (Fig. 3h) and performed co-IP experiments. 293T cells were co-transfected with expression constructs encoding HA-tagged Hoxa5 deletion constructs and V5-tagged Scip. Pull-down experiments with an antibody against the V5 epitope showed that HA-Hoxa5-ΔN86 and HA-Hoxa5-ΔN144 do not co-IP with Scip (Fig. 3i), suggesting aminoacids 30-86 at the N-terminal region of Hoxa5 are essential for complex formation with Scip.

To test whether the Hoxa5/Scip interaction can also be observed in vivo, we prepared whole tissue lysate from the cervical spinal cord of e12.5 mouse embryos and performed co-IP. Similar to transiently transfected 293T cells, we were able to IP Hoxa5 using a goat anti-Scip antibody. Further probing the blot with a rabbit-anti-Scip antibody, we were also able to detect Scip in the same IP fraction (Fig. 3j). However, we were unable to detect Hoxa5 or Scip in whole-cell lysate, likely due to lower endogenous expression.

To test if Hoxa5 and Scip expression is sufficient to suppress LMC identity, we co-electroporated constructs expressing mouse Hoxa5 and Scip under a pCAGGs promoter in chicken embryos, which lack phrenic MNs. We found that the overexpression of Hoxa5 and Scip did not affect the number of MNs generated, as electroporated cells still expressed normal levels of Isl1/2, but suppressed the expression of Foxp1 (Fig. 3k). Our data collectively indicate that Hoxa5 and Scip cooperate to induce phrenic and suppress limb-innervating MN identity.

### Postnatal maintenance of phrenic MN identity

Our data revealed mechanisms that control the establishment of embryonic phrenic MNs, largely through transcriptional programs mediated by Hox5 and Scip proteins. However, it is not clear whether expression of these two TFs is continuously required for phrenic MN maintenance at postnatal and adult stages. Since a number of Hox proteins show maintained expression at postnatal stages in brachial MNs^42^, we evaluated the expression of Hoxa5 and Scip at different stages. Both Hoxa5 and Scip were strongly expressed in phrenic MNs at postnatal day (P)5.5, but their expression became weaker at P10.5 and undetected by P16.5 (Fig. 4a).

**Figure 4.**
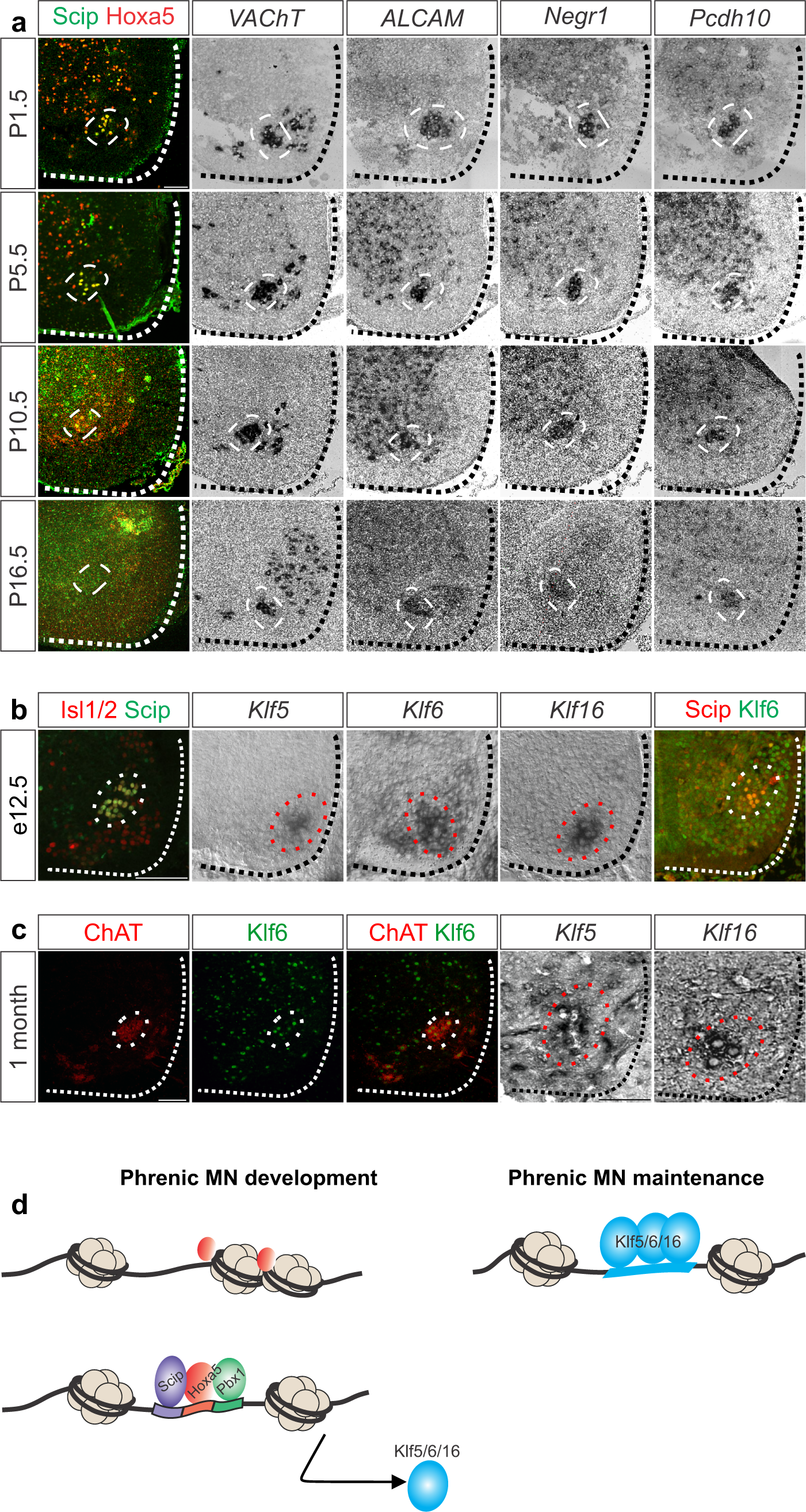
Maintenance of phrenic MN identity at postnatal stages. **a)** Expression of Hoxa5, Scip, *VAChT*, *ALCAM*, *Negr1* and *Pcdh10* in postnatal phrenic MNs. MNs are shown inside the dashed white line. Hoxa5 and Scip are downregulated after P10.5, while in situ hybridization shows sustained expression of phrenic-specific genes *Alcam*, *Negr1*, and *Pcdh10*. **b-c)** Expression of *Klf5*, *Klf6* and *Klf16* in phrenic MNs at e12.5 **(b)** and 1 month **(c)**. **d)** Proposed model of phrenic MN specification and maintenance. Hoxa5 can bind to inaccessible chromatin and forms a complex with Pbx1 and Scip to induce PMC-specific genes, including Klf factors, which may act to maintain phrenic MN properties in adulthood.

During development, Hox5 proteins control the expression of phrenic-specific cell adhesion molecules, such as *ALCAM, Negr1 and Pcdh10*^32^. To test whether Hox5 downstream genes are downregulated in a similar temporal fashion as Hoxa5 or are maintained postnatally, we performed in situ hybridization for a pan-MN marker *Vesicular Acetylcholine Transporter (VAchT)*, *Alcam, Negr1, and Pcdh10*. Surprisingly, we observed maintained expression of these genes at P16.5, despite Hoxa5 downregulation, suggesting that additional gene regulatory mechanisms may control the maintenance of these early Hox5 target genes (Fig. 4a). In order to explore potential maintenance factors of phrenic MN identity downstream of early Hox/Pbx programs, we intersected Hoxa5 and Pbx1-enriched ChIP-seq peaks and differential ATAC-seq peaks with a curated list of mouse TFs^43,44^. We selected several TFs that either showed particularly high enrichment in ChIP-seq or ATAC-seq datasets or have known functions in MN development for further downstream analysis by in situ hybridization. This analysis identified a number of TFs, including Ebf and Tshz factors, Neurod1, Onecut2, and Stat3. However, we did not observe phrenic-specific enrichment of these TFs at e12.5 (Fig. S3a). We also identified several Klf family members in our intersected dataset and previously noticed that the footprinting score of multiple Klf TFs was reduced in Hoxa5-deficient ATAC-seq peaks (Fig. 1i), suggesting Hoxa5 may regulate both the expression and DNA-binding of Klf family members. We tested Klf expression at e12.5 and found that Klf5, Klf6 and Klf16, but not Klf3, Klf7, or Klf15, are highly expressed in phrenic MNs (Fig. 4b, S3b). We also found that Klf5, Klf6, and Klf16 expression is maintained in phrenic MNs at 1 month of age (Fig. 4c). Together these findings suggest that a subset of Klf TFs show continuous expression from embryonic to postnatal phrenic MNs and may regulate a gene regulatory network required for phrenic MN identity maintenance (Fig. 4d).

## Discussion

Combinatorial TF expression and changes in chromatin accessibility underlie the development, diversification and maturation of MN subtypes^45^. Hox proteins are at the core of early transcriptional programs that diversify MNs along the rostrocaudal axis of the spinal cord^3^. At cervical levels of the spinal cord, MN columns show a differential requirement for Hox5 proteins-PMC neurons are largely dependent on Hox5 proteins for their survival and specification, LMC neurons show an intermediate requirement for the axonal pathfinding of a subset of pools, while MMC neurons appear to be resistant to Hox5 loss. Here, we sought to address how Hox5 proteins can serve multiple functions in the development and specification of distinct MN subtypes. We find that Hox5 paralogs exert their functions through altering chromatin states and associating with MN-specific co-factors. Our findings provide insights into how Hox5 proteins can selectively control both PMC and LMC properties. The high incidence of Hox motifs in open chromatin and Hoxa5 binding in MMC-associated genes is surprising, given the lack of overt MMC phenotypes in Hox5 mutants. While MMC columnar identity is thought to be Hox-independent, it is possible that Hox-mediated transcriptional programs may contribute to MMC properties downstream of columnar identity, similar to LMC neurons.

Several Hox proteins exert their functions partially through their ability to reorganize chromatin, a characteristic of pioneer factors^25^. We identify Hoxa5 as an additional family member that exhibits pioneer activity. It is unclear why Hox paralogs differ in their abilities to alter chromatin state. Despite substantial redundancy among Hox proteins in limb-innervating MN development, both Hoxa5 and Hoxc9 have unique abilities to induce phrenic and preganglionic/hypaxial MN identities, respectively^16,41^, and these distinct functions may partly arise from their ability to bind and open inaccessible chromatin, consistent with the idea that increased selectivity may be associated with lower chromatin accessibility^24,25,28^. Given the absence of domains that indicate an intrinsic ability of Hox proteins to remodel chromatin, it is likely that this property arises from their interactions with additional binding partners^25^. Both Oct and Klf family members have known pioneer activity^46^, indicating that the ability of Hoxa5 to recruit these TFs could mediate its chromatin remodeling activity.

Our data indicate that Hoxa5 has the differential ability to recruit Scip (Pou3f1/Oct6) and that this interaction is mediated by sequences at the N-terminal domain of the protein, which are the most divergent among Hox paralogs and thus likely to mediate paralog-specific protein interactions^47,48^. The ability of Hoxa5 to interact with this novel binding partner may have led to the emergence of phrenic MN identity in mammals, as avian species express Hox5, but not Scip, with similar rostrocaudal boundaries in the spinal cord. In mouse embryonic stem cell (ESC)-derived MNs, co-expression of Hoxa5 and Scip induces a transcriptional profile corresponding to phrenic MNs^19^. Here, we show that Hoxa5 and Scip co-expression is also sufficient to suppress LMC identity, revealing that the Hox5/Scip complex has a dual role in inducing phrenic and suppressing limb MN programs. Similarly, we previously found that combinatorial expression of Hoxa5 and FoxP1 suppresses phrenic MN identity^16^, indicating that cross-repressive interactions ensure the right balance of phrenic and limb-innervating MNs at cervical levels of the spinal cord. Interestingly, FoxP1 and Scip expression domains overlap at more caudal levels of the brachial spinal cord that are devoid of Hoxa5 expression, indicating that Hoxa5 is specifically required for Scip/FoxP1 cross-repression. Motif analysis of ATAC-seq and ChIP-seq data indicates a different top motif for Hoxa5 binding in PMC neurons, although this analysis is limited by the small number of known phrenic-specific genes. One possibility is that the interaction of Hoxa5 with Scip can bias its binding preferences to regulatory regions on phrenic-specific targets, suggesting conserved strategies for Hox binding selectivity^49–51^. Future experiments utilizing scRNA-seq, scATAC-seq and CUT&RUN from isolated phrenic MNs will further test this possibility.

The transcriptional programs that control MN maturation and maintenance are just beginning to emerge. In C. elegans MNs, terminal selectors are necessary for inducing and maintaining cholinergic transmission and other core features of MN identity throughout the lifetime of the animal^52–54^. In mammalian serotonergic neurons, an adult stage transcriptional program maintains their synaptic connectivity and protects axons from neurotoxic injury^55^. It is unclear whether mammalian MNs express maintenance factors that safeguard their integrity in adulthood, and whether these factors are broadly expressed in all MNs or are unique to specific MN subtypes. We find that a subset of Klf TFs are induced and maintained in phrenic MNs after downregulation of early Hox transcriptional programs, suggesting that they may act to maintain phrenic MN properties. Despite convergence of transcriptional programs in the majority of MN subtypes as they progress from development to adulthood, phrenic MNs appear to sustain their unique identity, as they form a distinct cluster in adult scRNA-seq data^56^. While Klf6 is broadly expressed in all adult MNs^45^, Klf5 and Klf16 expression appears to be more restricted, suggesting phrenic-specific functions. In addition to phrenic MNs, Klf5 is also expressed in hypaxial MNs in the thoracic spinal cord, indicating a conserved role in respiratory MN populations^19,57^. Klf family members differentially regulate the intrinsic ability of CNS axons to regenerate^58^, raising the possibility that specific family members may be broadly involved in neuroprotection or degeneration in adulthood.

## Methods

### Mouse genetics

The *loxP-*flanked *Hoxa5*^59^, *Olig2::Cre*^60^, and *ChAT(BAC)-eGFP* (*ChAT::GFP*)^61^ lines were generated as previously described and maintained on a mixed background. Mouse colony maintenance and handling was performed in compliance with protocols approved by the Institutional Animal Care Use Committee of Case Western Reserve University. Mice were housed in a 12-hour light/dark cycle in cages containing no more than five animals at a time.

### Immunohistochemistry and in situ hybridization

In situ hybridization and immunohistochemistry were performed as previously described^16,32^, on tissue fixed for 2 hours in 4% paraformaldehyde (PFA) and cryosectioned at 16μm. Postnatal mice (P5.5-P16.5) were perfused with a solution of phosphate-buffered saline (PBS) and 4% PFA, followed by a 2-hour post-fixation at 4°C. In situ probes were generated from e12.5 cervical spinal cord cDNA libraries using PCR primers with a T7 RNA polymerase promoter sequence at the 5’ end of the reverse primer. All probes generated were 750-1000bp in length. The following antibodies were used: guinea pig anti-Hoxa5, guinea pig anti-Hoxc5^33^, rabbit anti-Hoxb5^16^, guinea pig anti-FoxP1^22^, goat anti-Scip (1:5000; Santa Cruz Biotechnology, RRID:AB_2268536), mouse anti-Islet1/2 (1:1000, DSHB, RRID:AB_2314683), rabbit anti-Lhx3^62^, rabbit anti-Klf6 (1:1000, Santa Cruz Biotechnology, Cat# SC-7158) and goat anti-ChAT (1:200, Millipore, RRID:AB_2079751). Images were obtained with a Zeiss LSM 800 confocal microscope and analyzed with Zen Blue, ImageJ (Fiji), and Imaris (Bitplane).

### MN dissociation and fluorescence-activated cell sorting

C2-C6 cervical spinal cords were dissected from e12.5 embryos in a *ChAT::GFP* background in ice cold PBS and collected in PBS. After spinning down, the pellets were dissociated with Papain Dissociation System (Worthington, Cat# LK003176) following the manufacturer’s instructions. Briefly, tissue was enzyme digested for 30 min at 37°C with DNase (117 units/mL) and gently triturated. The single cell solution was centrifuged and then resuspended in PBS with 1% BSA and DNase. Dissociated cells were filtered through a 70 μm filter and subjected to fluorescence-activated cell sorting (FACS) on a BD Aria-SORP digital cell sorter with 85 μm nozzle to enrich for GFP positive cells. The cells were collected in a microtube containing 100 μL of PBS with 1% BSA.

### ATAC-seq library preparation

Bulk ATAC-seq for each condition was performed with at least two biological replicates as previously described^63^ and scaled down to half. Briefly, 25,000 FAC-sorted cells were centrifuged at 500 g for 6 min in a chilled centrifuge to form a pellet. The pellet was washed once in 25 μL of ice cold PBS, resuspended in 25 μL of cold lysis buffer (10 mM Tris-HCl, pH 7.4, 10 mM NaCl, 3 mM MgCl2, 0.1% Igepal CA-630) and centrifuged at 500 g for 10 min at 4°C. The cell pellet was resuspended in transposition reaction mix (12.5 μl TD-Buffer, 1.25 μl Tn5, 11.25 μl water) (Nextera DNA Library Prep Kit, Illumina, Cat# 15028212) and incubated for 30 min at 37°C. Immediately following the transposition reaction, purification was carried out using mini elute PCR Purification Kit (Qiagen, Cat# 27104). The appropriate number of amplification cycles was determined using qPCR reaction as described^63^. The PCR cycles were carried out with Illumina Nextera adapter primers using the NEBNext High Fidelity 2x Master Mix (NEB, Cat# M0541S) using the following PCR program: (1) 5 min at 72°C, (2) 30 s at 98°C, (3) 10 s at 98°C, (4) 30 s at 63°C, (5) 1 min at 72°C, and (6) repeat steps 3–5 with total cycles <12. Final PCR products were cleaned using PCRClean Dx beads (Aline Biosciences, Cat# C-1003) and assessed for quality using a Bioanalyzer. The libraries were sequenced on an Illumina NextSeq 550 (paired-end 75 bp) at the Genomics Core Facility at Case Western Reserve University.

### ATAC-seq data processing and analysis

ATAC-seq data were processed using the standardized uniform Encyclopedia of DNA Elements (ENCODE) pipeline from the ENCODE consortium^64^. Briefly, FastQC (v0.11.9) was used to check the pre-alignment read quality. FASTQ files from ATAC-seq reads were mapped to UCSC mm10 with Bowtie2 (v2.3.4.3). All unmapped reads, non-uniquely mapped reads, PCR duplicates and ChrM reads were removed using Samtools (v1.9). Peaks were called using MACS2 (v2.2.4) with parameters “--nomodel --shift 37 --ext 73 --pval 1e-2 -B --SPMR --call-summits”. Peaks overlapping with the blacklist region defined by ENCODE were removed using Bedtools (v2.29.0). Next, replicated peaks in each condition were intersected using Bedtools (intersect) to identify open chromatin regions overlapping by at least 1bp and defined as replicated peaks. Replicated peaks were annotated in R using the ChIPseeker package (v1.36.0), which assigns each peak to the nearest gene transcriptional start site (TSS). To identify differential peaks, FeatureCounts was used to obtain count data from the resulting ATAC-seq BAM files. Count data for all replicates and experimental conditions were combined into a single count matrix in R. The consensus peaks were identified as the peaks that were present in at least two samples. The count matrix was subsequently used to identify differentially expressed genes with the R package DEseq2^65^. PCA was performed using plotPCA function within DEseq2 on Variance Stabilizing Transformation (VST)-transformed data. Proximal and distal peaks were defined by associating differential ATAC-seq peak distances to annotated TSS (ChIPseeker). Peaks that were at least 2.0 kb away from the annotated TSS were assigned as distal ATAC-seq peaks, while all others were assigned as proximal. To visualize the ATAC-seq signal in the UCSC genome browser, samples were normalized to 1x genomic coverage, also known as Reads per Genome Coverage (RPGC).

### Motif analysis

HOMER (v4.10) was used to perform de novo motif enrichment^29^. Motif analysis on ChIP-seq data was performed using a fixed 200 bp window around the peak center. Motif analysis on ATAC-seq data was performed using a fixed 500 bp window around the peak center on differentially accessible chromatin. In both cases, the HOMER findMotifsGenome.pl command was used to perform de novo analysis against background sequences generated by HOMER that match the GC content. The top-scoring motifs, along with their p-value and enrichment, are shown.

### Footprinting analysis

To analyze footprinting signatures in ATAC-seq data the TOBIAS package^34^ was used. All replicates from each condition were merged into one .bam file using bedtools. Peaks were called using MACS2 with parameters “--nomodel --qvalue 0.01 --keep-dup all”. Peak files were associated with motifs from JASPAR CORE Vertebrates collection 2022^66^. Merged BAM files were processed using ATACorrect to correct for Tn5 bias. Footprint scores were calculated using FootprintScores, and differential footprinting analysis was performed using BINDetect.

### Go enrichment

The enrichGO function from the clusterProfiler (v4.8.2) package was used to perform GO term analysis of enriched biological processes and generate the graphs with maximum of 500 genes set for each category. The top ten significant GO terms were plotted and ordered by the number of gene counts in each category.

### Single cell RNA-sequencing (scRNA-seq) reanalysis

The filtered matrix output from the Cell Ranger pipeline for rostral samples was obtained from the Gene Expression Omnibus repository with accession code GSE183759^30^. Seurat package (v4.4.0) was used to perform quality filtering, normalization, dimensionality reduction, and cell clustering. Briefly, cells were evaluated for quality, and those with gene counts between 1000 and 5300, UMI counts below 30500, and mitochondrial counts under 10% were kept for further analysis. After filtering, 5460 cells were retained for downstream analysis. The resulting digital data matrices were then processed using a SCT transformation^67^ to perform normalization, scaling, and identification of variable features with mitochondrial reads regressed out. MNs were separated by the expression of common MN markers such as Mnx1 or cholinergic markers such as ChAT or Slc18a3 or Slc5a7. Only the cells expressing MN markers were considered for downstream analysis leading to a total of 5011 cells. To identify cell clusters, Uniform Manifold Approximation and Projection (UMAP) was used with the first 30 principle components. Cells were clustered using FindClusters function (resolution = 0.3) and visualized using UMAP. Cell identities were assigned using known markers. Clusters that were close to each other in UMAP space expressing LMC (*FoxP1*, *Aldh1a2*) and MMC (*Mecom*, *Lhx3*) markers were merged to create a new cluster ID and defined as LMC and MMC clusters. Furthermore, conserved markers for LMC and MMC clusters were generated by using Findconservedmarker function with logFC thresholds of 0.25. To identify ATAC-seq peaks associated with MN clusters in the cervical spinal cord, the conserved marker genes obtained from scRNA-seq for LMC and MMC were intersected with the genes associated with the ATAC-seq peaks.

### ChIP-sequencing (ChIP-seq)

e12.5 mouse cervical spinal cords were dissected and flash frozen in liquid nitrogen. The tissue samples, along with antibodies, rabbit anti-Hoxa5^33^ and rabbit anti-Pbx1 (Cell Signaling Technology, RRID:AB_2160295) were sent to Active Motif for chromatin isolation and sonication, ChIP assay, library preparation, library QC, Next-Generation sequencing on the Illumina platform and analysis. In brief, 75-nucleotide sequence reads generated by Illumina sequencing (NextSeq 500) were mapped to the mm10 genome using the BWA algorithm with default settings. Alignments that were uniquely mapped to the genome and had no more than two mismatches were retained for subsequent analysis. PCR duplicates were further removed. Peaks were called using MACS2 using the default p-value cutoff. Peak filtering was performed by removing ChIP-seq peaks aligned to the blacklist genome as defined by ENCODE. Peaks were annotated in R with the ChIPseeker package, which assigns each peak to the nearest gene’s TSS.

### Plasmid construction for co-immunoprecipitation (co-IP) and electroporation

To create expression vectors for co-IP experiments, RNA extracted from mouse spinal cord was converted to cDNA and used to amplify Hoxa5, Pbx1, and Scip using custom oligonucleotides with HA, Myc, and V5-tags. PCR amplified products and cloning vector (pcDNA3.1) were digested to create compatible sites for ligation and transformed into NEB10 beta competent bacteria (NEB, Cat# C3019H). To create plasmids for chick electroporation, mouse Hoxa5 and Scip cloned into the pcDNA3.1 vector were used to amplify Hoxa5 and Scip and inserted into pCAG-tdTomato (Addgene, Cat #83029), a vector with the chick β-actin promoter/CMV enhancer. The complete length of cloned plasmids was sequenced at Eurofins and verified by mapping to the respective mRNAs using the UCSC mouse reference genome.

### Co-immunoprecipitation (co-IP) assays

HEK293 cells were transfected using Lipofectamine 3000 (Invitrogen, Cat# L3000008) according to the manufacturer’s instructions. After 48 hours, cells were washed once in ice cold PBS and harvested in 1X RIPA buffer (Cell Signaling, Cat# 9806). Co-IP assay was carried out using protein A/G PLUS-Agarose beads (Santa Cruz, Cat# 2003). Briefly, 600 μg of total cell lysate was precleared with 20 μl of agarose beads for 30 min. For co-IP, 200 μg of precleared protein was incubated with 2 μg of anti-HA (Cell Signaling Technology, Cat# 5017, RRID:AB_10693385, fig. 3b-e), anti-V5 (Santa Cruz Biotechnology, Cat# sc-271944, RRID:AB_10650278, fig. 3g, 3i) or anti-Myc (Cell Signaling Technology, Cat# 2276, RRID:AB_331783, fig. 3f) and incubated for 1 hour on a rocker at 4°C. To conjugate beads with the antibodies bound to the protein, 20 μl of agarose beads were added and incubated at 4°C overnight. Protein complex bound beads were washed 3 times with RIPA and 2 times with PBS and the pellet was resuspended in 40 μl of 1x sample buffer and boiled for 3 minutes. 25 μL of the immunoprecipitated aliquots and 5% of total lysate (input control) were run on a standard SDS-PAGE gel. The gels were then transferred onto a PVDF membrane (BioRad, Cat# 1620177) using a wet transfer system and blocked by incubation with 3% BSA in TBST (TBS with 0.1% Tween-20). Membranes were probed with anti-HA, anti-V5, anti-Myc or anti-Scip (Santa Cruz Biotechnology, RRID:AB_2268536). Blotted membranes were scanned using Odyssey infrared imaging system (Li-COR).

For in vivo co-IP, cervical tissue from e12.5 mouse embryos was washed once in ice cold PBS and homogenized in RIPA buffer (60μL/embryo) using a disposable pestle. The lysate was incubated at 4°C for 30 min and then clarified by spinning down at 4°C for 10 min at 10,000 RPM. 200 μg of precleared lysate was incubated with 2 μg of goat anti-Scip (Santa Cruz Biotechnology, RRID:AB_2268536) and incubated for 1 hour on a rocker at 4°C. To conjugate beads with the antibodies bound to the protein, 20 μl of agarose beads were added and incubated at 4°C overnight. Protein complex bound beads were washed 5 times in PBS and the pellet was resuspended in 40 μl of 1x sample buffer and boiled for 3 minutes. 20% of total lysate was used as input control for running a standard SDS-PAGE western blot. After transfer, the blot was blocked and probed with rabbit anti-Hoxa5, washed, and re-probed with rabbit anti-Scip (RRID:AB_2631304).

### In Ovo Electroporation

Electroporation was performed by introducing a DNA solution into the lumen of the neural tube of specific pathogen-free (SPF) chicken embryos (AVS Bio, Cat#10100326) at Hamburger-Hamilton stages 14-16^68^ using 5× 50 msec pulses at 25V, with electrodes placed horizontally across the longitudinal axis of the embryo to achieve unilateral electroporation of the desired construct mixture. The DNA solution was composed of the relative ratios of each construct diluted in TE buffer with 0.5% Fast Green to aid injection visualization. The construct concentrations were adjusted to obtain a final ratio of 2:2:1 for Hoxa5:Scip:EGFP in which the total DNA electroporated per egg was 1.1µg/µl. Electroporated embryos were incubated at 37°C for 3 days and analyzed at stages 25-26.

### Statistics and reproducibility

The programs used for data analysis such as MACS2 for peak calling, DEseq2 for differential analysis, Homer for motif-enrichment analysis, clusterprofiler for GO term enrichment analysis, and Tobias for footprinting score analysis use algorithms that provide their own p values, q values, and/or FDR. The data was primarily analyzed in R (v 4.3.1) and the R scripts used for data analysis are freely available upon request. For electroporation experiments, data are represented in violin plots to show overall frequency distribution of all the individual data points, with dashed lines representing the median value and dotted lines representing the two quartile lines. P-values were calculated using a one-way ANOVA. p < 0.05 was considered to be statistically significant, where * p< 0.05.

## Data availability

All sequencing data produced for this study will be available at the Gene Expression Omnibus (GEO) upon publication.

## Code availability

The R scripts used for data analysis are freely available from the corresponding author, PP, upon request.

## Acknowledgements

We thank Heather Broihier, Evan Deneris, Britton Sauerbrei, Ashleigh Schaffer and Helen Miranda for discussions and comments on the manuscript. We thank Clay Spencer and Evan Deneris for assistance and helpful discussions on bioinformatics analysis, Ayana Sawai, Jeremy Dasen and Lynn Landmesser for in ovo-electroporation protocols, Jeremy Dasen for Hox5, FoxP1 and Lhx3 antibodies and Ganapati Mahabaleshwar for the Klf6 antibody. We also thank the Cytometry and Imaging core at Case Comprehensive Cancer Center (P30CA043703) for assistance with FACS and Active Motif for assistance performing ChIP-sequencing. This work was funded by NIH R01NS114510 to PP, F31NS120699 and T32GM008056 to RKC. PP is the Weidenthal Family Designated Professor in Career Development.

## Author contributions

RKC and PP conceived and designed the study, RKC, RLB, ML and PP performed experiments and analyzed data, LJ provided *Hoxa5 floxed* mice, RKC, RLB and PP wrote the paper with input from all authors.

---

**Figure S1.**
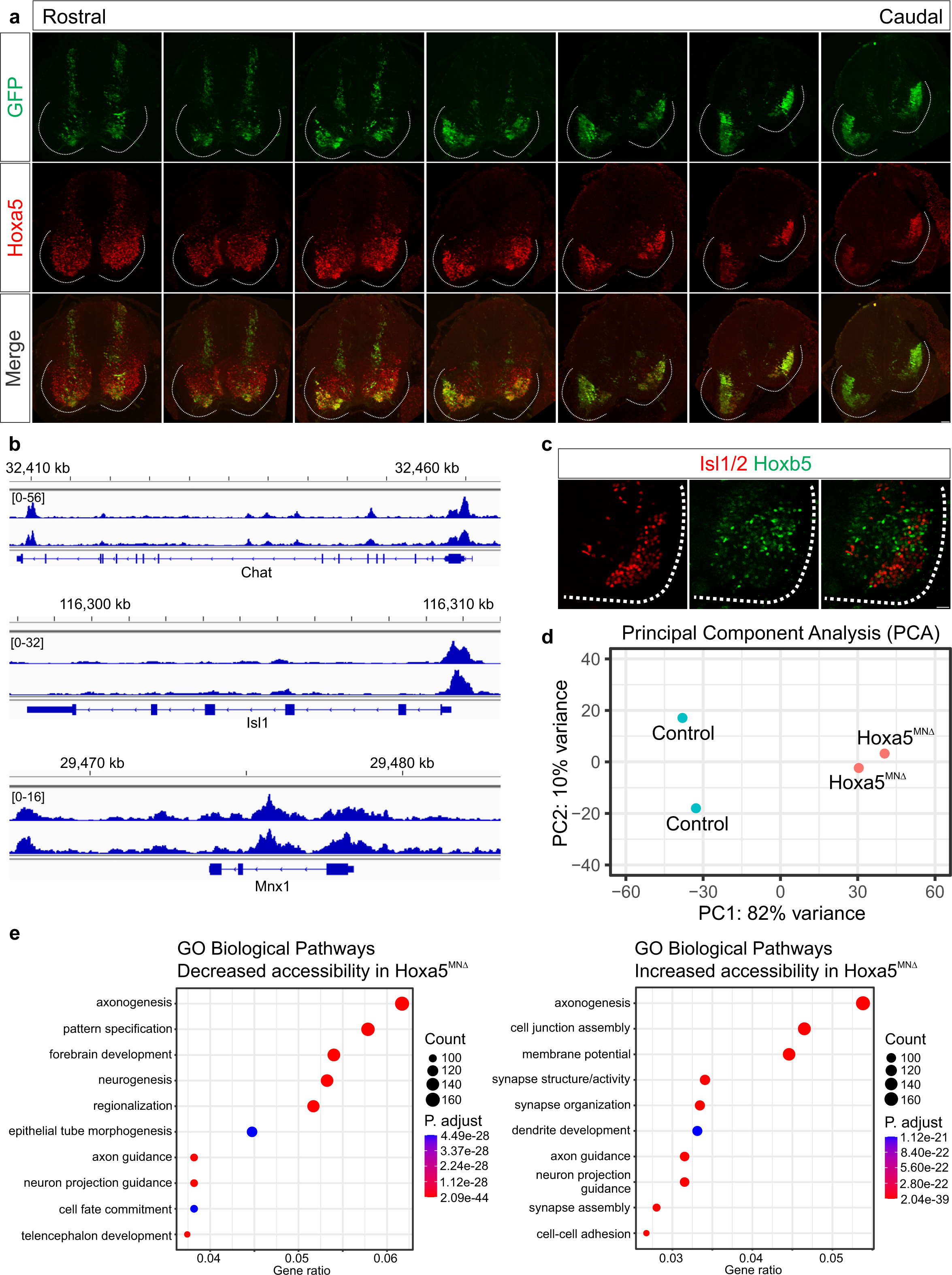
Hoxa5 contributes to chromatin accessibility in cervical MN subtypes. **a)** Rostral to caudal distribution of e12.5 *ChAT::GFP+* MNs (green) at spinal cervical levels, within the boundaries of Hoxa5 expression (red), sorted for ATAC-seq analysis. **b)** Genome browser views of ATAC-seq signals from two *ChAT::GFP+* control samples at three MN-specific genes: *ChAT*, *Isl1* and *Mnx1*. **c)** Hoxb5 (green) is not expressed in MNs (Isl1/2+ in red) at cervical spinal cord levels. **d)** Principal component analysis (PCA) for control and Hoxa5-deleted MNs reveals changes in the accessible chromatin landscape of cervical MNs after Hoxa5 deletion. **e)** Gene ontology (GO) enrichment analysis of biological pathways of genes with changes in chromatin accessibility in *Hoxa5^MNΔ^* MNs. Top 10 significant GO enrichment pathways shown based on the gene counts in each category. Scale bar= 50μm.

**Figure S2.**
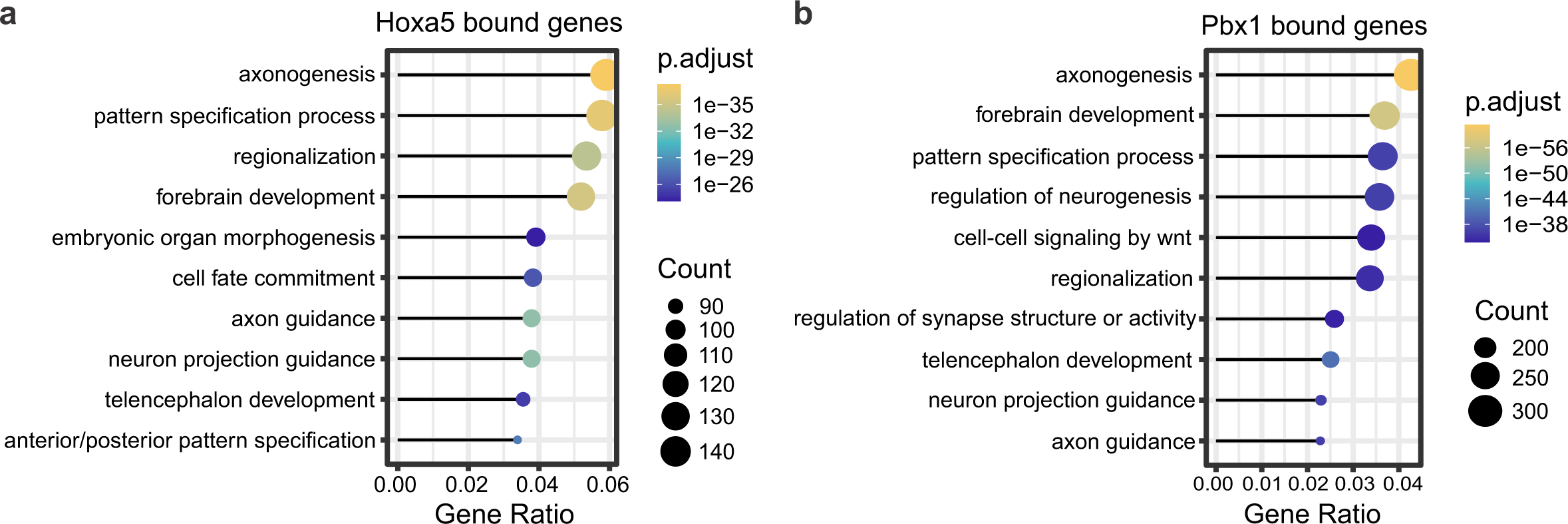
Direct regulation of MN-specific genes by a Hoxa5/Pbx1 complex. **a-b)** Gene ontology (GO) enrichment analysis of biological pathways of genes with Hoxa5 **(a)** and Pbx1 **(b)** ChIP-seq peaks. Top 10 significant GO enrichment pathways shown based on the gene counts in each category.

**Figure S3.**
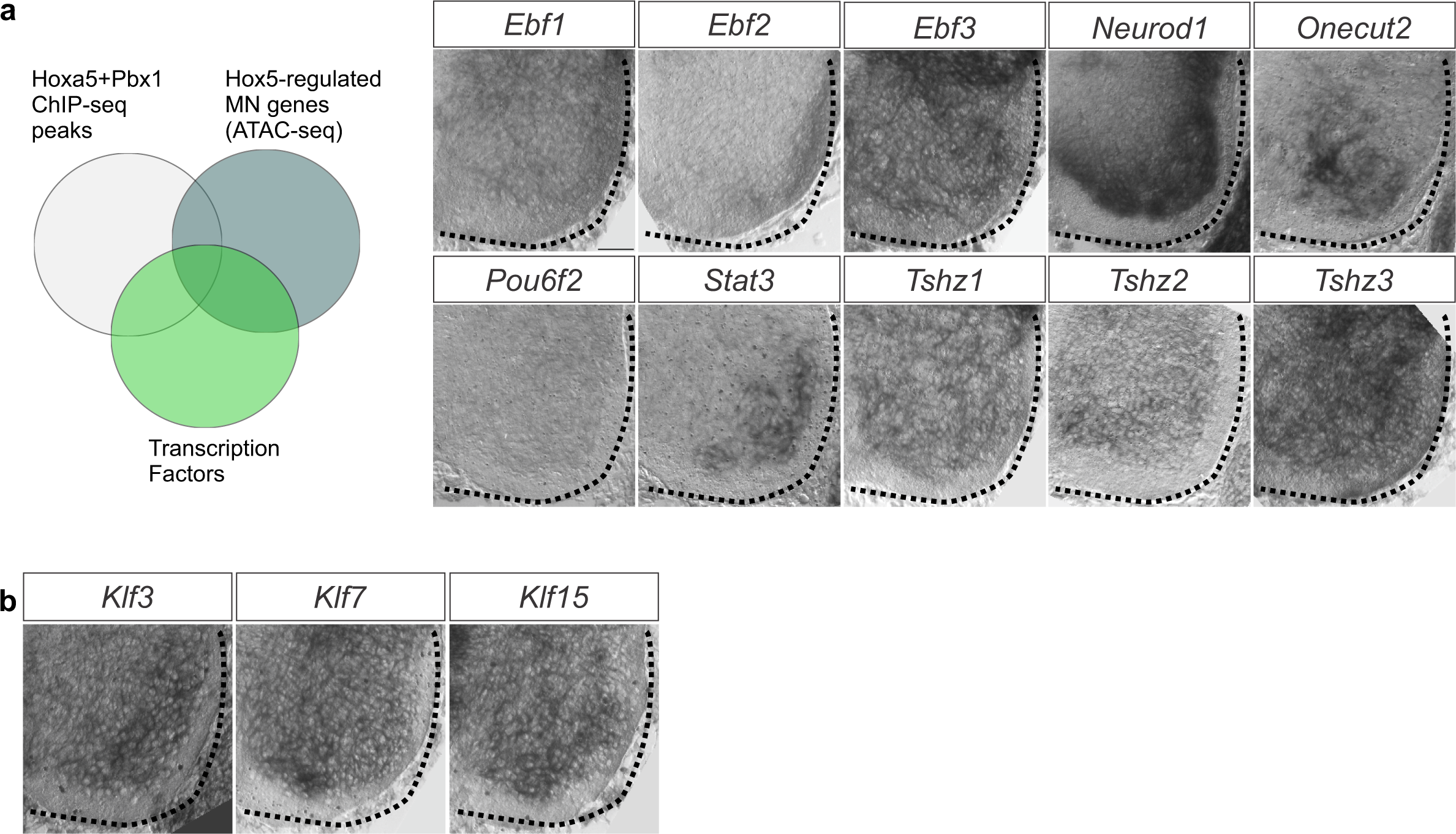
A transcription factor code for phrenic MN maintenance. **a)** In order to identify transcription factors acting downstream of Hoxa5 and Scip in phrenic MNs, we intersected Hoxa5 and Pbx1-enriched ChIP-seq peaks, differential ATAC-seq peaks between control and *Hoxa5^MNΔ^* MNs and a transcription factor dataset. Several TFs were identified but did not show phrenic-specific expression by in situ hybridization. **b)** *Klf3*, *klf7* and klf15 were not enriched in phrenic MNs at e12.5. Scale bar= 50μm.

**Table S1.** Column-specific MN genes with ATAC-seq peaks. Intersection of scRNA-seq data with ATAC-seq data to identify accessible chromatin regions in LMC- and MMC-expressed genes. For PMC peaks we used a list of previously identified PMC-specific genes.

| LMC-enriched genes |  |  | MMC-enriched genes |  |  | PMC-enriched genes |
| --- | --- | --- | --- | --- | --- | --- |
| Snhg6 | Cadm2 | Cplx1 | Kcnb2 | Dhrs1 | Pcdh7 | Cdh9 |
| Epha4 | Pcp4 | Kcnd2 | Khdrbs2 | Kctd12 | Nwd2 | Cdh10 |
| Cdh7 | Epb41l3 | Tspan12 | Igfbp5 | Lifr | Apbb2 | Pcdh11x |
| Lypd1 | Rit2 | Ptn | Ecel1 | Laptm4b | Slc10a4 | Pcdh10 |
| Atp2b4 | Syt4 | Tmem176b | Adora1 | Nell2 | Adgrl3 | Ptn |
| Brinp3 | Onecut2 | Npy | Lhx4 | Copz1 | Epha5 | Hoxa5 |
| Rgs4 | Grp | Skap2 | Mgst3 | Uts2b | Cplx1 | Hoxa1 |
| Tagln2 | Tszh1 | Snca | Rxrg | Lsamp | Sdk1 | Alcam |
| Akap12 | Scd2 | Reep1 | Atp2b1 | Epha3 | Rasl11a | Negr1 |
| Mrpl54 | Sorcs1 | FoxP1 | Tmtc2 | Robo2 | Ubl3 | Plxnc1 |
| Socs2 | C1ql3 | Ret | Lin7a | Ncam2 | Snx10 | Vwc2l |
| Nudt4 | Lrp1b | D930028M14Rik | Rab3ip | Ezr | Hoxa9 | Vwc2 |
| Csrp2 | Stk39 | Fxyd7 | Grip1 | Clic1 | Creb5 | Edil3 |
| Nefh | Itga6 | Ldha | Bcl11a | Ppp2r2b | Cyp26b1 | Mmd2 |
| Vstm2a | Ppp1r1c | Nr2f2 | Kcnip1 | Sema6a | Rassf4 | Synpr |
| Pank3 | Map1a | Rgs10 | Ebf1 | Vim | Cd9 | Pappa |
| Lhx1 | Gatm | Inpp5f | Rasgef1c | Nacc2 | Ntf3 | Hs6st2 |
| Lhx1os | Ctxn2 | Ifitm2 | Sar1b | Lhx3 | Nell1 | Klf5 |
| Nme2 | Tpd52 | Cend1 | Sez6 | Neurod1 | Slco3a1 | Zbtb7c |
| Etv4 | Anxa5 | Zdhhc2 | Bcas3 | Lrrc4c | Cbln1 | Lsamp |
| Rac3 | Pcdh10 | Mt3 | Ngfr | Mpped2 | Crnde | Ptprt |
| Hpcal1 | Trim2 | Tppp3 | Pitpnc1 | Scg5 | Irx5 |  |
| Dtnbp1 | Crabp2 | Nrp1 | Sox9 | Myt1 | Maf |  |
| Enc1 | S100a11 | Ntm | Churc1 | Mecom | Gria4 |  |
| Cartpt | Camk2d | Anxa2 | Cplx2 | Sertm1 | Aplp2 |  |
| Pnoc | Negr1 | Aldh1a2 | Ube2ql1 | Fbxw7 | Aw551984 |  |
| Lgals1 | Dab1 | Polr2m | Med10 | Rhoc | Cadm1 |  |
| Prph | Hpca | Clstn2 | Irx1 | Lmo4 | Megf11 |  |
| Hoxc5 | Ywhah | Tenm1 | Gm17750 | Ddah1 | Foxb1 |  |
| Hoxc4 | Atp8a1 | Arhgap36 | Pik3r1 | Epha7 | Prss35 |  |
| Smug1 | Gabrg1 | Pcdh19 | Rgs7bp | Brinp1 | Gria3 |  |
| Abat |  |  | Il6st | Ptprd | Hprt |  |
|  |  |  | Isl1 | Camk2n1 | Fgf13 |  |
|  |  |  | Rarb | Casz1 | Sema3c |  |
|  |  |  | Slc18a3 |  |  |  |

